# piNET: a versatile web platform for downstream analysis and visualization of proteomics data

**DOI:** 10.1101/607432

**Authors:** Behrouz Shamsaei, Szymon Chojnacki, Marcin Pilarczyk, Mehdi Najafabadi, Chuming Chen, Karen Ross, Andrea Matlock, Jeremy Muhlich, Somchai Chutipongtanate, Dusica Vidovic, Vagisha Sharma, Juozas Vasiliauskas, Jake Jaffe, Michael MacCoss, Cathy Wu, Ajay Pillai, Avi Ma’ayan, Stephan Schurer, Mario Medvedovic, Jarek Meller

## Abstract

Large proteomics data, including those generated by mass spectrometry, are being generated to characterize biological systems at the protein level. Computational methods and tools to identify and quantify peptides, proteins and post-translational modifications (PTMs) that are captured in modern mass spectrometers have matured over the years. On the other hand, tools for downstream analysis, interpretation and visualization of proteomics data sets, in particular those involving PTMs, require further improvement and integration to accelerate scientific discovery and maximize the impact of proteomics studies by connecting them better with biological knowledge across not only proteomics, but also other Omics domains. With the goal of addressing these challenges, the piNET server has been developed as a versatile web platform to facilitate mapping, annotation, analysis and visualization of peptide, PTM, and protein level quantitative data generated by either targeted, shotgun or other proteomics approaches. Building on our experience with large scale analysis of gene and protein expression profiles as part of the Library of Integrated Network Cellular Signatures (LINCS) project, piNET has been designed as a fast, versatile and easy to use web-based tool with three modules that provide mapping from peptides (with PTMs) to proteins, from PTM sites to modifying enzymes that target those sites, and finally from proteins (with PTMs) to pathways, and for further mechanistic insights to LINCS signatures of chemical and genetic perturbations. piNET is freely available at http://www.pinet-server.org.

## INTRODUCTION

Methods and tools for computational proteomics data analysis evolve constantly in order to match rapid advances in proteomics, and enabled by them large scale efforts in proteomic profiling of biological systems (1–3). In this context, many methods and data processing tools have been developed for the identification and quantification of peptides and proteins from biological samples using mass spectrometry-based approaches (4–6). Further downstream analysis typically follows to facilitate biological interpretation of such obtained quantitative results, with its own distinct set of challenges (6), and an interdependent ecosystem of databases, tools and resources for proteomics research (7).

Increasingly, proteomic profiling efforts involve identification and quantification of specific proteoforms that may exhibit distinct activities and functions, including those resulting from protein post-translational modifications (8). The importance of protein phosphorylation, acetylation, methylation and other PTMs that are involved in essential cellular processes has led to the development of databases and tailored resources, such as PhosphositePlus (9), Phospho.ELM (10), and PhosphoPep (11) for phospho-proteomics, as well as comprehensive databases of PTMs and aggregators related functional annotations, such as iPTMnet (12), dbPTM (13) or SIGNOR (14).

This contribution introduces a web server for both interactive and programmatic downstream analysis and visualization of peptide and protein level proteomic data, with special emphasis on PTMs. The new tool, dubbed piNET, aims to streamline mapping, annotation and analysis of post-translational modification sites, as well as modifying enzymes that target these sites, in the context of biological pathways. As illustrated in Figure 1, piNET integrates iPTMnet (Protein Information Resource), PhosphositePlus, SIGNOR, and other proteomic resources with pathway and PTM network analysis, while making use of a fast peptide mapping and versatile visualization engines.

**Figure 1.**
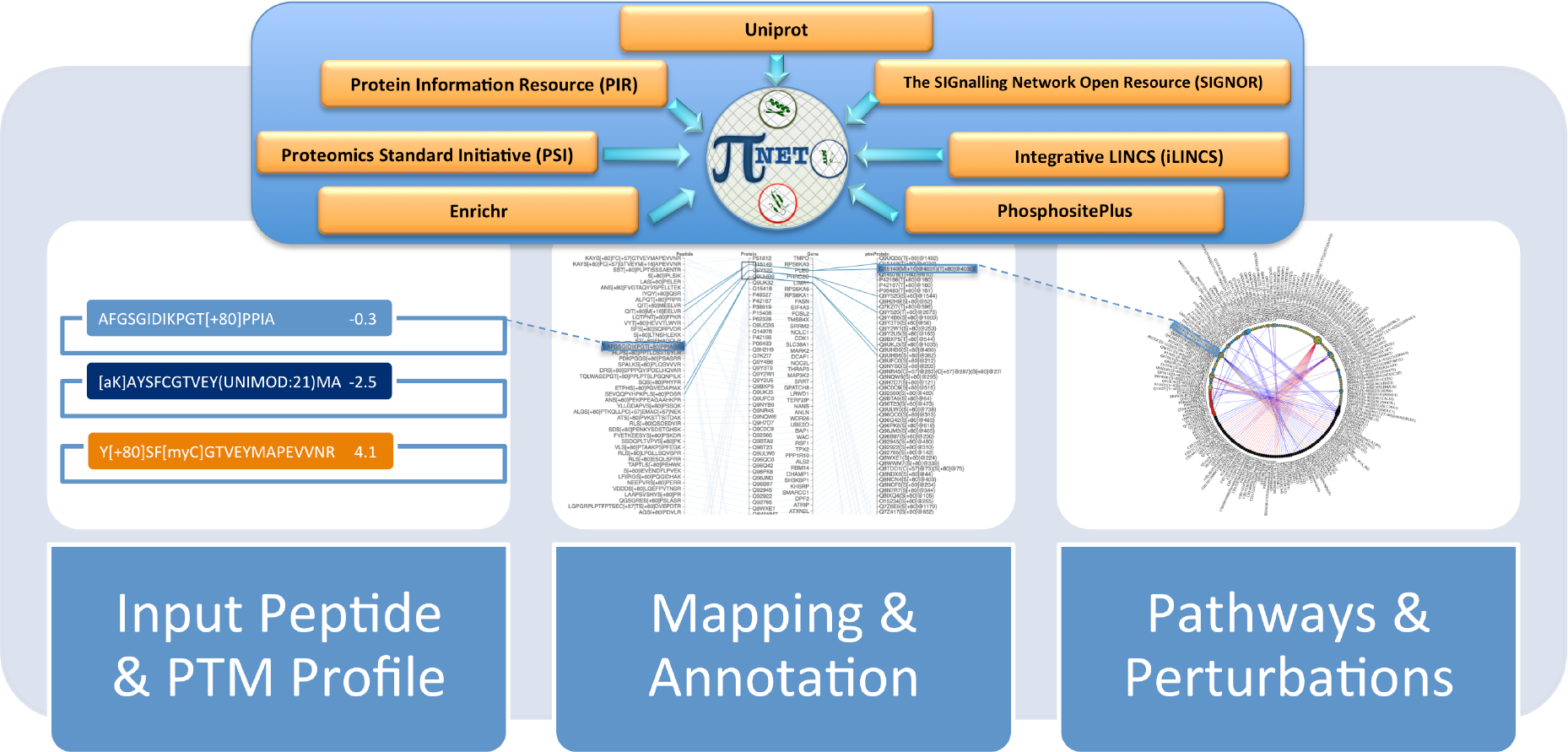
The overall flowchart of piNET mapping, annotation and visualization workflow.

Furthermore, piNET connects proteomics profiles with transcriptional and proteomics signatures generated within The Library of Integrated Network-based Cellular Signatures (LINCS) project (15), which aims to systematically collect Omics signatures of genetic and chemical perturbations, including those for the modifying enzymes, such as kinases, induced either by their genetic knock-downs or small molecule inhibitors. By utilizing seamless integration with Enrichr (16–17), iLINCS (http://ilincs.org) and related tools that facilitate interaction with the LINCS library of signatures, one can apply the Connectivity Map approach (18) in order to gain further mechanistic insights into signalling cascades that may be driving the underlying biological states and their proteomics profiles.

## MATERIAL AND METHODS

piNET has been implemented as a modular web server, using the Spring-Boot JAVA platform in conjunction with state-of-the-art D3 JavaScript libraries (and their in-house developed extensions) for high quality visualization. As a result, piNET can be executed within the browser and does not require installing additional plugins or applications for analysis and visualization. Programmatic access to piNET workflows is enabled through the corresponding RESTful API methods. Examples of Python scripts that invoke these respective APIs and return the results in the json format are included, to illustrate how they can be embedded into other applications. Additional documentation is provided in the ‘About’ pages to illustrate piNET workflows and the use of APIs with realistic examples.

piNET integrates a number of databases and resources for proteomics studies (see Table 1) in order to streamline mapping, annotation and analysis of post-translational modification sites, as well as modifying enzymes that target these sites. Multiple definitions of PTMs can be used interchangeably, including shorthand notation (e.g., pS, aK), mass difference (DeltaMass) identified experimentally, or PTM ontologies, including PsiMOD (https://www.ebi.ac.uk/ols/ontologies/mod) and UniMOD (http://www.unimod.org). As part of the initial processing of input peptide/PTM lists, using a Pride-MOD utility (19) and in-house developed translation modules, PTMs are mapped into PsiMOD and the resulting unified encoding can be used (in conjunction with mapping into proteins) as a basis for proteoform identifiers (provisionally implemented in the form of Protein Line Notation tokens).

**Table 1.**
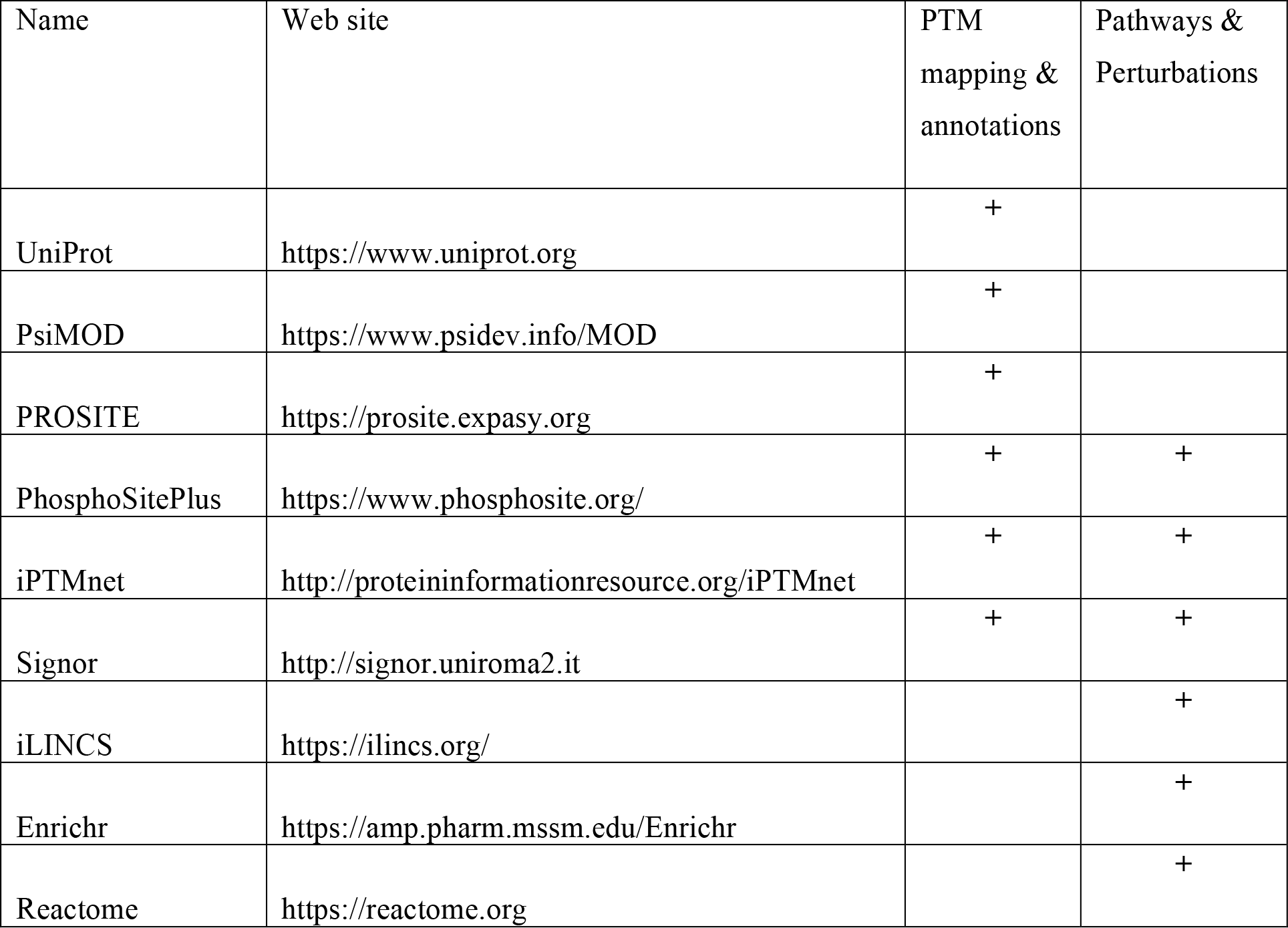
The list of databases and resources integrated in piNET with the goal of providing an easy to use and intuitive interface for the analysis and visualization of proteomics data.

piNET can be used to query Uniprot or Prosite APIs in order to map peptides and PTM sites into proteins from well annotated and continuously updated proteomes available through these authoritative resources (20–22). However, for fast mapping of large sets of peptides and PTMs, a locally indexed version of the UniProtKB/Swiss-Prot and TrEMBL protein sequence databases for three commonly used organisms in the proteomics profiling, i.e. Homo Sapiens, Mus Musculus and Rattus Norvegicus, has also been implemented using the state-of-the-art Apache Lucene indexing by Chen et al. (23). Using this fast peptide matching approach, an average search time of 0.02 seconds per peptide has been observed in our tests for a random sample of 20,000 peptides of length 10.

In addition, in order to overcome the limitations on the total number of peptides that can be submitted directly due to the maximum length of the URL, a file upload mode can be used to submit large sets of peptides and PTMs (with their abundance values). This option also enables submission of datasets with proteomic profiles of multiple samples and meta-data descriptors to define groups of samples, with the goal of deriving a simple differential expression profile for further analysis. Volcano plots can be generated in this workflow to guide the choice of differentially expressed peptides/PTMs/proteins based on the analysis of the effect size (fold change) and statistical significance (t-test based p-values).

Furthermore, piNET mines iPTMnet, PhosphositePlus, and SIGNOR for functional annotation of PTMs, and integrates these annotations to identify kinases and other modifying enzymes that are experimentally known to target PTM sites involved, and as a basis for PTM network analysis and visualization. For kinases, experimentally known modifier-PTM site pairs are augmented by identifying sites that share high degree of sequence motif similarity with known target sites (using PhosphoSitePlus known target sequences and consensus motif definition), and thus are likely phosphorylated by the same kinases. piNET, with its modular annotation and visualization engines can be easily extended to include representative, well benchmarked methods, such as NetworkKIN (http://networkin.info/index.shtml) or AKID (http://akid.bio.uniroma2.it) for the prediction of kinase-substrate pairs and other related resources (27–29) to further expand these putative annotations.

Finally, piNET connects protein level profiles with transcriptional and proteomics signatures generated by the LINCS project (30). piNET utilizes the aforementioned Enrichr APIs to generate the results of enrichment analysis, with emphasis on gene sets consisting of gene up- and/or down-regulated in response to kinase loss or gain of function, an thus representing kinase gain or loss of function signatures. The enrichment results are integrated with protein (gene) level expression profile visualization in order to connect further likely modifying enzymes and their down- or up-regulated targets. In a similar vein, piNET uses iLINCS APIs to facilitate further analysis of protein (or gene) level concordance with LINCS signatures, especially with those induced by the loss or gain of function of the modifying enzymes.

## RESULTS

We believe that piNET provides an intuitive and easy to use interface, divided into modular workflows. Nevertheless, a number of use cases, including those stemming from the LINCS project, are incorporated in order to illustrate its potential advantages, network-based visualization and integration with biological domain knowledge. We are also using these use case to illustrate potential pitfalls, such as those related to the problem of projecting proteoform level data onto gene level analysis.

### Peptide to Protein Workflow

As the first step in downstream analysis, piNET can be used to annotate, map and analyze a set of peptide moieties, including those with post-translationally modifications. Starting from a set of peptides, and optional modifications within those peptides, piNET provides mapping into both canonical and other isoforms included either in the UniProtKB/Swiss-Prot and/or TrEMBL databases. piNET also provides mapping and harmonization of PTM meta-data, using PsiMOD and UniMOD ontologies, and generating Protein Line Notation tokens to represent each peptide moiety with modifications for automated annotation exchange systems. Mapping of peptides and PTM sites into proteins and genes is summarized as intuitive interactive graphs and tables that can be downloaded, and also generated using API calls for integration with other resources. An example of a graphical summary of peptide to protein to PTM mapping is shown in Figure 2 for the set of P100 phospho-peptides (32) that are being used to profile responses to cellular perturbation by the LINCS project.

**Figure 2.**
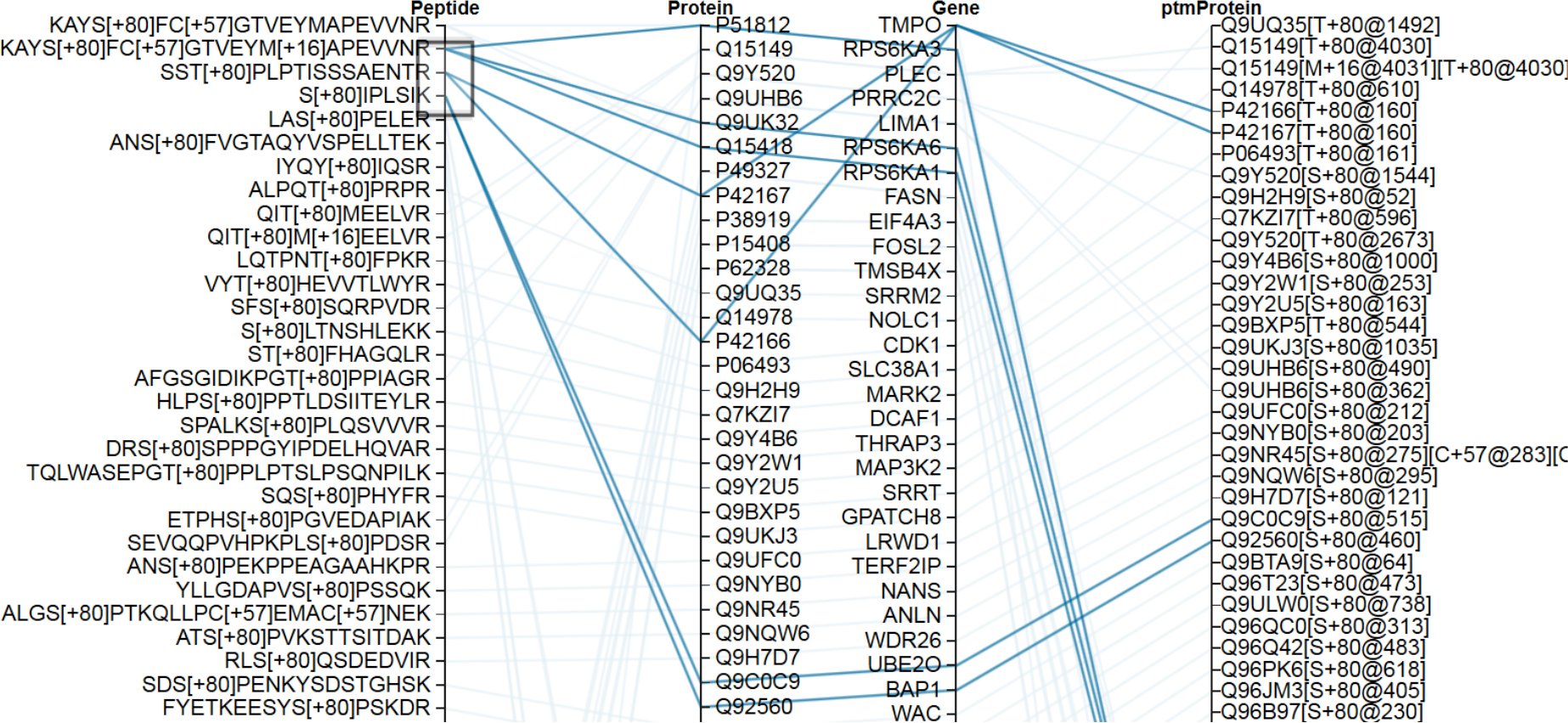
Peptide to protein to gene mapping for a subset of P100 representative phospho-peptides with peptides and PTM modification sites mapped into the corresponding protein and gene entities. Note that some peptides match multiple proteins and/or genes (many to many mapping), as indicated by thick blue edges for the 3 highlighted peptides.

### PTM to Modifying Enzyme Workflow

This PTM-centric workflow aims to elucidate signaling cascades converging on PTM sites, by mapping known and predicted upstream enzymes that target those PTM sites. Interactive high resolution PTM-centric network view is provided by embedding PhosphoSitePlus, iPTMnet and SIGNOR databases of enzyme-substrate relationships. For phosphorylation, known kinase - target peptides (and extrapolations from known examples to highly similar putative target peptides) are derived from PhosphoSitePlus (9) to provide a site specific PTM network view, and shed light onto phosphorylation cascades that may be involved. On the other hand, iPTMnet provides comprehensive annotation of not only kinase – PTM sites (which largely overlaps with that of PhosphoSitePlus), but also other modifying enzymes and their targets (12). Finally, SIGNOR can be used in piNET to map carefully curated causal relationships, available for over 2800 human proteins participating in signal transduction. Moreover, PTMs causing a change in protein concentration or activity have been curated and linked to the modifying enzymes (14).

The enzyme-substrate relations can be visualized in interactive graphs that help biological relations in the protein expression values. For example, the casual relationships extracted from SIGNOR can be represented as edges in the graph, indicating the activation/inactivation relationships between signalling entities by the color of an edge. A circular PTM network representation for P100 using SIGNOR causal relationships is illustrated in Figure 3. Several visualization options are available, including a bipartite graph, with the moieties that are being quantified in a proteomic assay shown as nodes in the right column, and modifying enzymes that target these PTMs in the left column. In each case, abundance levels of peptide/PTM/protein moieties can represented by the color of nodes, with yellow corresponding to high relative abundance and blue to low relative abundance, respectively. Such views provide a visual representation of relationships between quantified peptide/PTM/protein moieties, and other relevant biological entities or processes.

**Figure 3.**
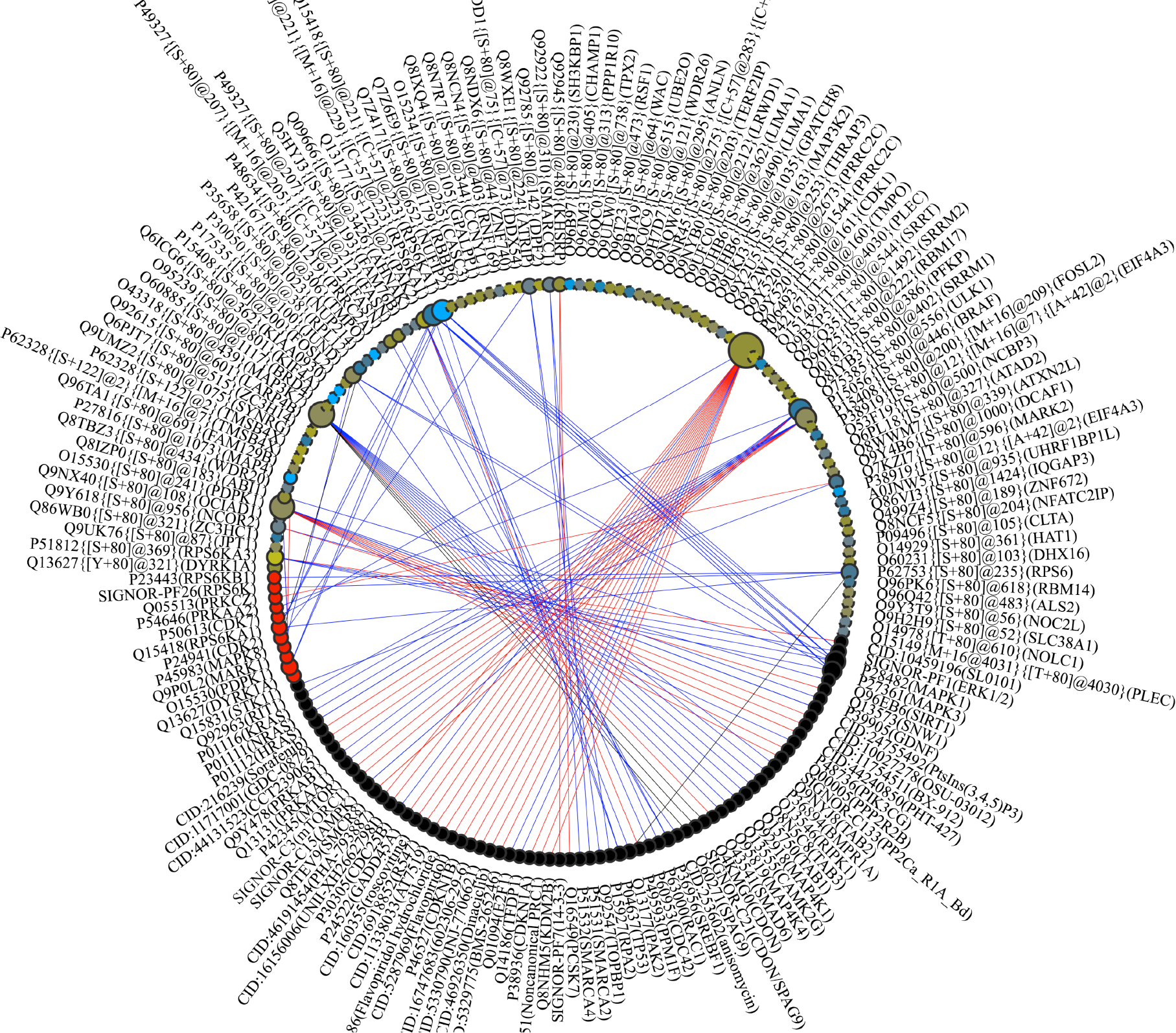
Functional annotation and visualization of a PTM network for a set of P100 phospho-peptide probes using curated causal annotations from SIGNOR, overlaid with a representative P100 expression (abundance) profile for a non-specific kinase inhibitor staurosporin generated as part of the LINCS project. Note that abundance of P100 peptides is indicated by the color of nodes representing them: blue for low and yellow for high abundance, respectively. Note also that some of the P100 phosphosites are connected via red or blue edges for positive or activating vs. negative or inactivating relationships, respectively, with targeting them modifying enzymes (red nodes), or other phosphorylation dependent modifiers, such as protein-protein interactions that depend on phosphorylation of the target site (black nodes).

### Protein to Pathway & Perturbation Workflow

A set of peptides (with PTMs) with the associated expression (abundance) values can be projected onto the protein (and thus gene) level for the purpose of further pathway analysis and visualization. The user can also defined per protein (gene) values when submitting protein level profiles in the Protein2Pathway tab. Subsequently, piNET can be used to facilitate pathway enrichment analysis with Enrichr, and to perform further mapping into Reactome pathways for interactive exploration, while generating high resolution visualization and tabulated results.

Since PTMs mediate cell growth, death and differentiation through intracellular signal transduction cascades and the resulting transcriptional changes, gene expression signatures can potentially also be used to capture biological states in the absence of direct proteomics profiles in response to molecular, genetic and disease perturbations, including the loss of function (due to genetic KDs and small molecule inhibitors) of modifying enzymes. To that end, piNET can be used to analyze protein level data in conjunction with LINCS signatures of cellular perturbations, by using iLINCS APIs or exploring the results interactively in iLINCS (http://ilincs.org).

## DISCUSSION

piNET has been designed to provide an integrated web platform for analysis, interpretation and visualization of large-scale proteomics data. To that end, piNET enables fast peptide/PTM to protein mapping, harmonization of meta-data pertaining to PTMs, systematic PTM to modifying enzyme mapping and other functional annotations, coupled with high quality visualization of PTM networks and protein pathways. While there are a number of tools and resources that provide much of the aforementioned functionality, including those listed in Table 1, we believe that piNET adds significantly to downstream proteomic data analysis by integrating these individual components and annotations resources, and by coupling them with a high quality visualization engine. In addition, to the best of our knowledge, there are no web-based tools that enable fast, large-scale mapping of peptides and PTMs, integrated with subsequent analysis of PTM networks for biological insights.

Furthermore, by virtue of mapping peptides, PTMs and their expression levels into proteins and genes, piNET facilitates connecting proteomics profiles with transcriptional and proteomics signatures generated by the LINCS project through the use of iLINCS and Enrichr as APIs. The goal is to provide additional mechanistic insights that may be revealed by connecting likely modifying enzymes with their down- or up-regulated targets through connectivity analysis (18). However, several inherent limitations of this approach must also be considered. In particular, mapping peptide and protein moieties from user defined profiles into the available LINCS targeted proteomics profiles such as P100 or GCP, may result in a small number of matches. Similar problems concern the use of L1000 transcriptional signatures, with additional caveats pertaining to their use as proxies for proteomic signatures that may not be available.

In this regard, we would like to comment that the LINCS data generation effort is on-going and we expect the number of signatures available as part of this step of analysis to grow over time. We also aim to further integrate piNET and its visualization engine with the recently published PTMsigDB that enables PTM-Signature Enrichment Analysis (PTM-SEA) of phospho signature data (32), and thus should alleviate some of the above mentioned limitations as well.

## ACKNOWLEDGEMENT

We would like to thank all of our early users, especially Drs. Rob McCullumsmith, Ken Greis, Michael Wagner and their groups, members of the BD2K-LINCS Data Coordination and Integration Center, as well as LINCS consortium participants, for their feedback and encouragement.

